# When Firing Rate Falls Short: Spike Synchrony Reliably Disentangles Stimulus Saliency and Familiarity

**DOI:** 10.1101/2025.08.09.669457

**Authors:** Viktoria Zemliak, Gordon Pipa, Pascal Nieters

## Abstract

Whether neural computation relies on firing rate or spike timing has been debated for decades, with no definitive resolution. Here, using a recurrent spiking network simulation and realistically varying stimulus saliency, we demonstrate that the two mechanisms are complementary: rate reliably detects stimulus class, spike synchrony reliably detects stimulus familiarity. This division of labor is necessary: rate coding fails to distinguish novel high-saliency from familiar low-saliency stimuli, while synchrony succeeds. We validate this complementary coding across a biologically realistic V1 model and an abstract associative memory network, demonstrating robustness across connectivity regimes and memory loads. Crucially, when inputs are already temporally coordinated through familiarity in a previous network layer, spike synchrony alone can encode both stimulus identity and familiarity. Our findings reconcile the rate-timing debate: familiar stimuli exploit efficient synchrony-based codes reinforced by recurrent connectivity across the cortical hierarchy, while novel stimuli depend on rate codes.

## Introduction

Is the firing frequency of action potentials sufficient to understand how neurons compute and communicate, or is the timing of individual spikes crucially important? Rate codes offer compelling advantages: they reliably correlate with experimental stimuli and behavior [1-3], are the basis of numerous theoretical and data-driven mean-field and plasticity analyses [4-6], and approximate successful deep learning networks [7]. Yet these advantages come at a cost: individual spikes carry little information, making rate codes inefficient and potentially inadequate to account for behavior on fast timescales [8-9]. Timing codes achieve high efficiency by encoding information in the precise order of spikes [10], time-to-first-spike in a stimulus response [11], rhythmic oscillations [12], bursting patterns [13], and spike synchrony [14]. While spike train variability challenges the reliability of timing codes [15], this variability may reflect meaningful recurrent interactions [16] or be mitigated by the relative nature of timing codes [17]. Thus, whether and to what extent brains rely on timing or rate information remains a fundamental yet unresolved question in neuroscience.

Rather than viewing rate and timing as competing alternatives, we propose they encode distinct, complementary information about stimuli. We designed a stimulus detection task with two key challenges: encoding stimulus identity across different saliency levels and distinguishing familiar from unfamiliar stimuli. Here we focus on absolute familiarity, whether stimuli match established patterns in recurrent connectivity, rather than relative familiarity from recent exposure, as these involve distinct neural mechanisms [18]. Two critical cases are particularly difficult but important to distinguish: highly familiar but weak stimuli and unfamiliar, but highly salient stimuli. A faint rustle from a known predator or a loud, unknown sound might evoke similar firing rates yet require fundamentally different responses. While sensory representations are generally believed to be encoded in firing rates [1-3], recent evidence suggests that absolute familiarity manifests as spike synchrony emerging from strong recurrent connections [19-21]. This suggests a complementary scheme where rate and spike-time coding coexist in the same network, each carrying distinct information about the stimulus.

We tested this complementary coding hypothesis using two recurrent spiking networks: a biologically realistic V1-like model and an abstract associative memory network. Across both models, synchrony reliably detected stimulus familiarity despite varying input saliency – a task where rate coding failed. Conversely, firing rates accurately identified which neurons encoded the stimulus, regardless of familiarity. This division of labor proved robust across connectivity regimes and was most pronounced in energy-efficient, sparse firing regimes. When familiar stimuli induce synchronized activity, this synchrony could propagate through a network, enabling pattern classification using synchrony. Our findings suggest that rate and timing codes work together, potentially reconciling the longstanding debate between these coding schemes.

## Results

### Experimental setup

We examined how neural circuits simultaneously encode stimulus identity and familiarity under realistic conditions of varying stimulus saliency. Stimulus saliency and familiarity are two kinds of meta-information about incoming stimuli, in addition to information about the specific content of perceptions, such as which objects are depicted on an image. They provide higher-level context about its relevance for processing and decision making.

**Saliency** reflects bottom-up sensory characteristics of stimuli, such as contrast and intensity. This distinguishes saliency from relevance or priority, which involve top-down attention modulation [22]. Visual contrast and intensity influence firing rates across various brain areas, particularly in thalamus which provides feedforward input to visual cortex [23], analogously to feedforward input in our simulations. Thus, stimulus saliency modeled as input firing rate varies significantly in our experiments, simulating real-world conditions.

**Familiarity** indicates whether a stimulus matches stored memories from prior experience. We focus on absolute familiarity linked to stable lifelong memories, rather than relative familiarity memory which reflects recent exposure [24]. Familiarity memory is essential for adaptive behavior and decision-making in both living and artificial agents, facilitating the recollection of well-known stimuli or, vice versa, the encoding of new stimuli in memory [24-25]. Lifelong absolute familiarity memory was suggested to be encoded in recurrent (lateral) connections in computational models [19-21]. One example is V1, where strong lateral connections are formed between neurons with similar receptive fields and in closer proximity to each other [26-27], which aligns with Gestalt-like statistics of natural visual stimuli [28-29]. Another prominent example is the hippocampal region CA3, where pyramidal neurons form ensembles which strengthen with experience and participate in associative memory recall [30]. Consequently, recurrent connections reflect the acquired experience and influence the level of spike synchrony [31-32].

Neural networks processing stimuli must simultaneously identify what each stimulus is, determine stimulus familiarity, and do so robustly across varying levels of stimulus saliency. In our experiments, we therefore varied saliency and familiarity in a stimulus detection task (Fig. 1). While output firing rate responds to both stimulus dimensions, spike synchrony can selectively reflect input familiarity [19]. This selectivity becomes crucial when saliency varies: firing rate alone cannot reliably distinguish between new but highly salient stimuli (Fig. 1B) and familiar but weak stimuli (Fig. 1C), as both can produce similar rate responses. In contrast, spike synchrony reliably differentiates these cases because synchrony is highly sensitive to recurrent connections that have been strengthened by repeated exposure to familiar stimuli [20-21]. Thus, we hypothesized that spike-time coding via synchrony can provide complementary information to rate coding.

**Figure 1.**
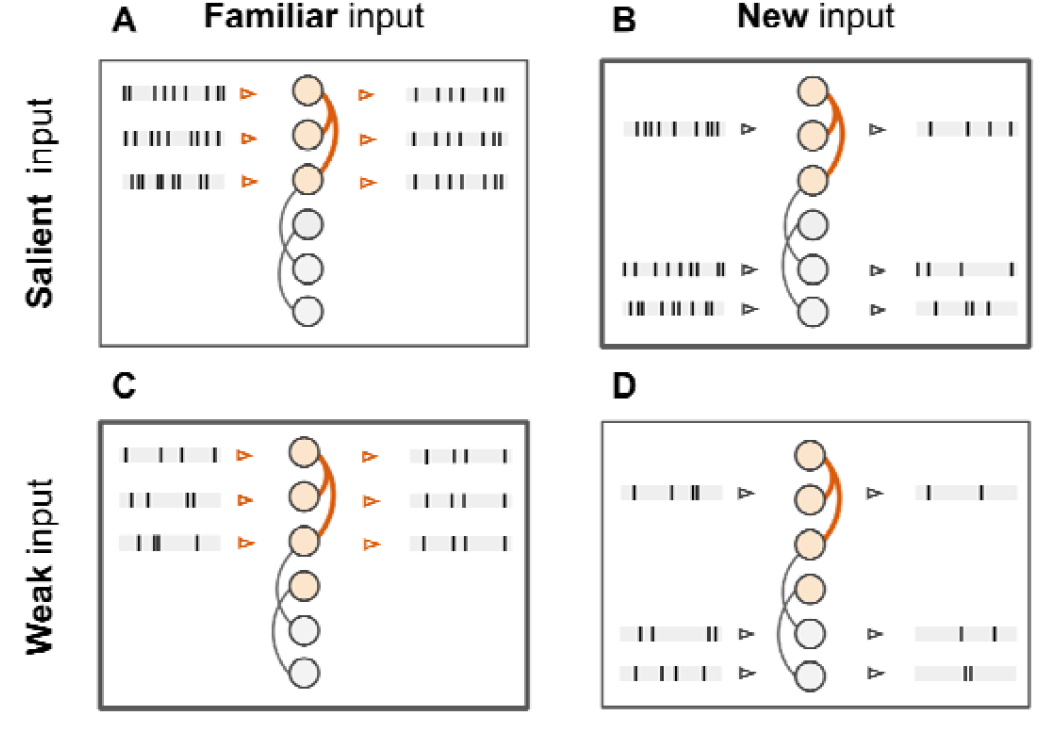
Distinguishing *salient new* inputs from weak *familiar* inputs is difficult. Input stimuli can be divided into 4 groups based on their meta-information: **A**. Familiar and salient, **B**. New and salient, **C**. Familiar but not salient (weak), **D**. New and not salient. Saliency is represented b the input firing rate, and familiarity by strength of recurrent connectivity. The difficult case to distinguish i between conditions B and C: When saliency varies, output firing rate (spike count) is similar between both, once driven by higher recurrent excitation, once by higher external excitation.. However, we hypothesize that they can be distinguished reliably by spike synchrony.

To test our hypothesis, we conducted a series of experiments in which recurrent spiking networks had to detect input familiarity and, in some cases, identify the subset of neurons receiving external stimulation (see Fig. 2B and the next two sections for experimental details). To predict familiarity, we used the network’s spiking activity in response to a stimulus sample. We trained a univariate logistic regression classifier for binary classification (1 = familiar, 0 = new), using either firing rate or spike synchrony as predictors. Firing rate was quantified as the average spike count (SC), and spike synchrony as Rsync (see Methods).

**Figure 2.**
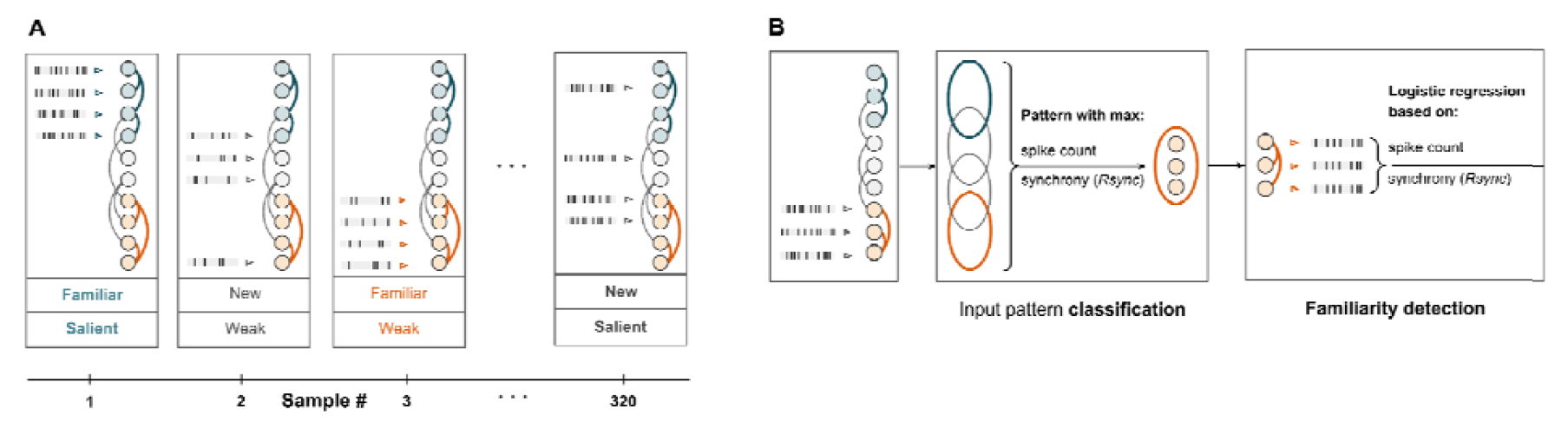
Familiarity detection and pattern classification on a dataset of new and familiar patterns with varying saliency. Dataset & experimental pipeline. In the network diagrams, teal and orange colors represent strongly connected clusters which encode familiar patterns.. **A**. Experimental procedure for both network models (V1 and associative memory model): 160 new and 160 familiar stimuli in random order, with varied saliency (input rate). Familiar patterns corresponded to network clusters, new stimuli did not.**B**. The abstract model of associative memory encodes multiple familiar stimuli. It first has to identify which neurons receive the feedforward input and then use this group of neurons to detect pattern familiarity.

For each sample in the simulations, external input was delivered to a subset of neurons − weakly interconnected for new stimuli and strongly interconnected for familiar ones − in a form of Poisson spike trains via one-to-one feedforward connections. These trains were asynchronous, uncorrelated, and maintained a constant rate within a sample. The input rate itself was drawn from a uniform distribution, with the range defining the input rate variability. Thus, in both experiments, input saliency (rate) varied across samples, for both new and familiar stimuli. Each dataset comprised 320 samples: 160 familiar and 160 new (see Fig. 2A).

### Familiarity detection under increasing input variability in a V1-type network

First, we tested our hypothesis in a 800-neurons recurrent spiking network that modeled connectivity of excitatory horizontal connections in V1, with connections and weight distribution based on experimental observations (see Methods). Familiar input patterns in the network were represented by clusters of 40 spatially neighboring neurons with similarly tuned receptive fields and strong mutual connectivity, consistent with cluster analysis of neurons in L2/3 of V1 [33-34]. This reflects the core principle of Hebbian plasticity in V1: neurons that are frequently co-activated tend to develop stronger synaptic connections over time which reflects acquired visual experience [35-36]. We assumed that prior experience with repeatedly occurring visual patterns lead to the emergence of tightly connected neuronal clusters. Therefore, we considered inputs matching one cluster of tightly connected neurons as familiar, whereas we considered patterns that innervated neurons across clusters as new inputs.

#### Input familiarity detection is more robust using spike synchrony

We measured how classification accuracy in familiarity detection depended on measuring spike synchrony and spike count in either: (global) all neurons in the model, or (local) the subset of neurons receiving external feedforward input (Fig. 3A). Both features perform better when evaluated for the group of neurons receiving the feedforward stimulus input rather than globally for an entire network.

**Figure 3.**
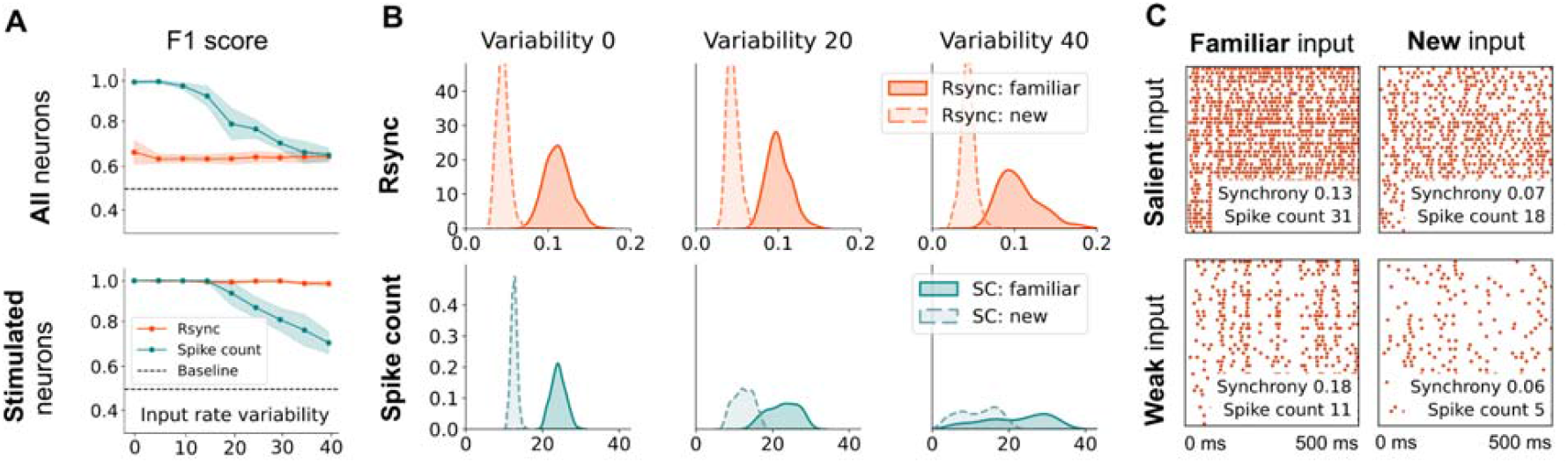
Familiarity detection for V1 model at different saliency (input rate) variability shows advantage for local synchrony-based decoding. Cross-validated performance of Rsync- and spike count-based classifiers for detecting input familiarity from all neuron activity and from the activity of externally stimulated neurons, averaged over 10 trials. Every trial includes 360 data samples, input saliency (firing rate) variability changed from 0 (50 Hz) to 40 (10-90 Hz). **A**. Averaged F1 score for different input saliency (firing rate) variability. Rsync-based familiarity detection outperforms spike count for higher rate variability. **B**. Kernel-density-estimation plots of synchrony and spike count of stimulated neurons in response to new and familiar stimuli. Rsync shows a clear separation across levels of saliency, whereas distributions for spike count start overlapping heavily. **C**. Example spike raster plots of 500 ms of network activity in response to weak (10 Hz input rate) and salient (90 Hz input rate), new and familiar stimuli, with spike synchrony and average population spike count for every condition.

For spike count this difference was minor, while synchrony benefits significantly from a localized measurement. Local synchrony-based familiarity classification performance remains impressively robust under conditions of significant saliency variability (up to 40 Hz) which strongly supports the role of spike timing in encoding input familiarity.

#### Distributions of spike synchrony are well-separated in all conditions

We also examined the distributions of spike synchrony and spike count among stimulated neurons (Fig. 3B). Spike synchrony distributions remained well-separated across all input variability conditions, while spike count distributions became indistinguishable under higher input variability. This demonstrates that spike synchrony is a reliable indicator of familiarity in dynamic environments: firing activity becomes more temporally coherent upon encountering a familiar stimulus (Fig. 3C).

### Pattern classification and familiarity detection under increasing memory load

To investigate the computational potential of a complementary rate and spike synchrony coding scheme, we conducted a familiarity detection experiment under the conditions of input rate variability 40 Hz emulating high variability in stimulus saliency (for details see Methods). Here, a 5000-neuron SNN model followed more abstract connectivity principles: We generated *n* familiar and *n* new binary input patterns, each defined by a subset of 100 externally stimulated neurons. For familiar patterns, each neuron formed strong excitatory connections with other neurons from the same pattern with probability 0.4, thus creating clustered representations. The quick emergence of such clusters and their use for familiarity recognition has been demonstrated in rate [37], excitatory spiking [21], and balanced E-I [38] networks. We additionally implemented cross-pattern inhibitory connections similar to associative memory networks [39-40]. The probability of inhibitory connections was 0.02. All connection weights were randomly drawn from the same log-normal weight distributions as in the V1 experiment for consistency. In sum, each neuron formed outgoing excitatory connections within its cluster and inhibitory connections with neurons from the other clusters. The resulting network performed two tasks:

1) Input pattern classification. This was achieved by selecting an input pattern with the highest SC or Rsync out of *n* familiar + *n* new patterns.

2) Familiarity detection of the pattern identified from SC (Fig. 4).

**Figure 4.**
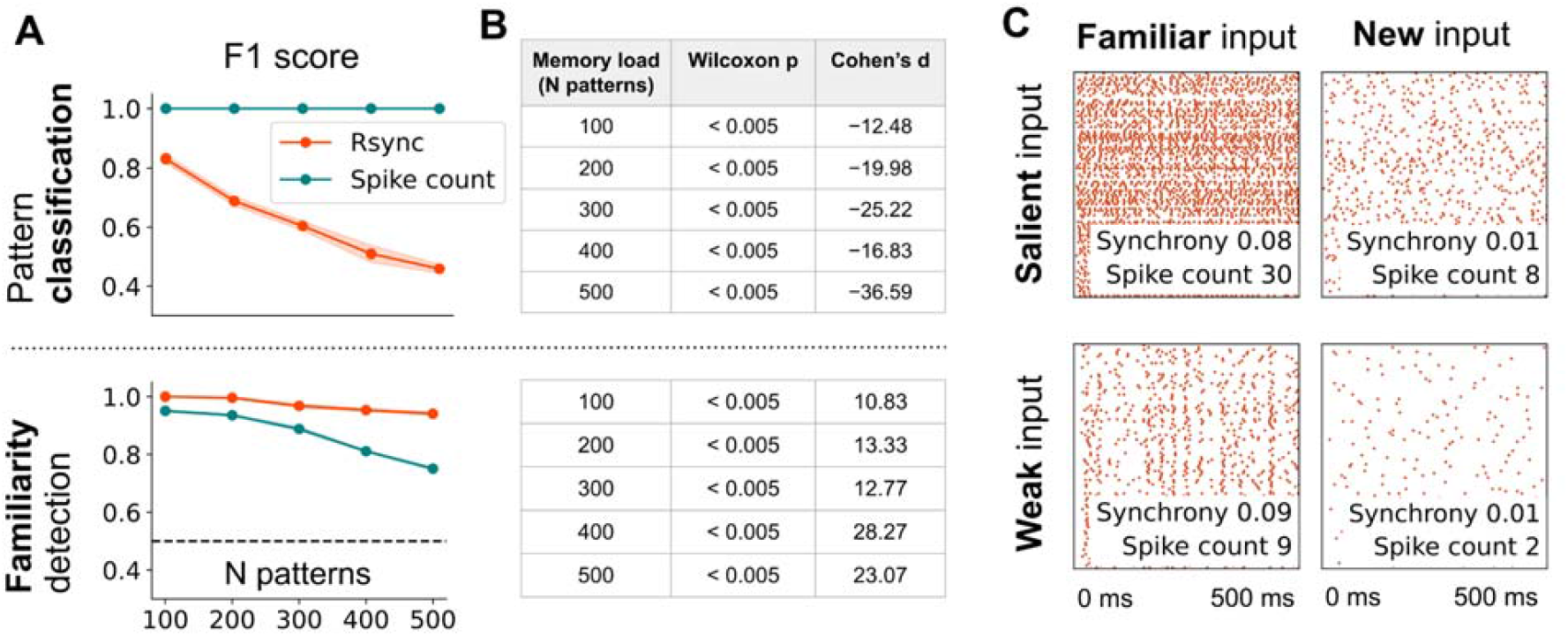
Familiarity detection and pattern classification in the associative memory model shows distinct roles for spike count and synchrony. Cross-validated performance of Rsync- and spike count-based classifiers for detecting a stimulated group of neurons (input pattern class) and the familiarity of a pattern which they are encoding, averaged over 10 trials. Every trial includes 360 data samples, input saliency (firing rate) variability across samples was 40 Hz. **A**. Averaged F1 score for different amounts of pre-encoded patterns and **B**. Wilcoxon test for significant differences between Rsync and spike count and Cohen’s d for the effect size demonstrate that, across memory load (number of encoded patterns in the network) Rsync outperforms spike count in familiarity detection, but falls behind in pattern classification. This strongly supports a complementary coding scheme using Rsync for familiarity and SC for pattern classification. **C**. Spike raster plots of 500 ms of network activity under memory load 100 in response to weak (10 Hz input rate) and salient (90 Hz input rate), new and familiar stimuli, with synchrony and average population spike count.

#### Rate reliably detects stimulus class, synchrony reliably detects familiarity

Nearly all new and familiar stimulated patterns were detected with 100% accuracy using average spike count, enabling familiarity detection from identified neurons. The Wilcoxon test showed significant superiority of spike count over Rsync for the classification of a stimulated pattern (Fig. 5B). This is particularly true for novel stimuli, which are exceedingly difficult to classify from spike synchrony. In contrast to classification, spike synchrony performed significantly better in a familiarity detection task under the maximum input saliency (input rate) variability 40 Hz (input rate range 10-90 Hz). The effect size measured as Cohen’s d was the strongest for the most challenging experiment condition: 500 pre-encoded familiar patterns (Fig. 4B). However, Rsync has an important limitation in terms of familiarity classification: it strongly relies on accurate pattern classification. Spike count and spike synchrony thus provide complementary information about the stimulus — its content and familiarity, respectively.

**Fig 5.**
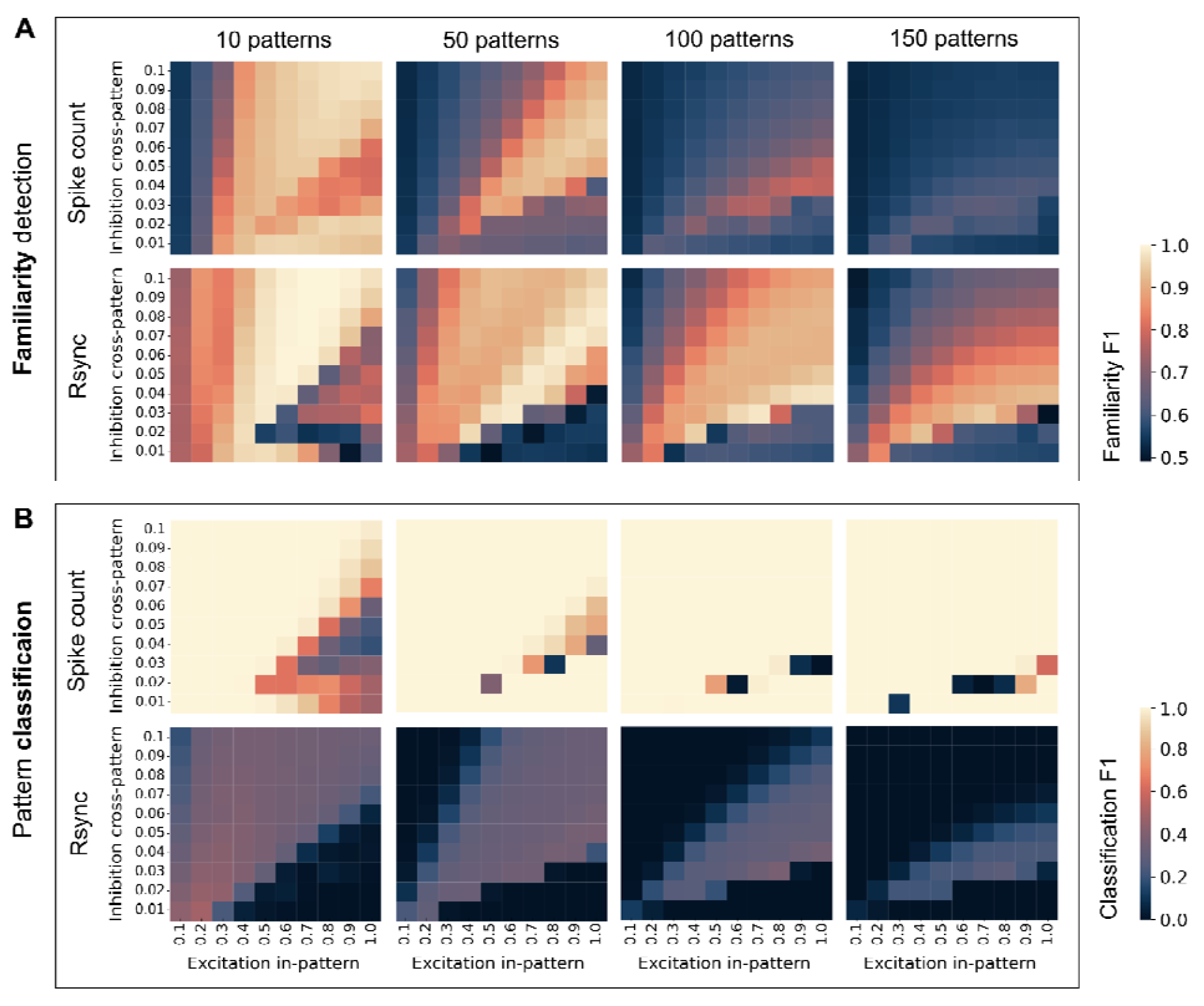
Familiarity detection and pattern classification across connectivity parameters. Cross-validated performance of Rsync- and spike count-based classifiers, for every combination of the proportion of in-pattern excitatory connections and cross-pattern inhibitory connections. Averaged over 10 trials, every trial includes 360 data samples, input saliency (firing rate) variability across samples 40 Hz. **A**. shows F1-scores for familiarity detection, **B**. shows macro F1-scores in pattern classification. Spike count performs better at pattern classification, while synchrony excels at familiarity detection, especially for more encoded patterns. Correspondingly, familiarity detection with Rsync is robust across multiple connectivity regimes and pattern classification with spike count is highly robust across connectivit regimes.

Both familiarity detection metrics show declining performance as the number of familiar patterns stored in the network increases. This effect aligns with the results of Zemliak et al. [21], who attribute it to growing overlap between clusters representing familiar patterns. In the next section, we show how this effect can be counteracted by regulating the proportion of excitatory and inhibitory connections.

### Rate and synchrony codes across different connectivity regimes

To ensure that the advantage of spike synchrony in familiarity detection was not due to a specific parameter setting, we systematically varied the proportions of excitatory and inhibitory connections. We compared the performance of rate-based and synchrony-based familiarity and pattern classification across a broad range of connectivity regimes and for different numbers of familiar patterns stored in networks of 1000 neurons. Weight amplitudes and distributions consistent with previous simulations. The experimental parameter space was defined by two variables: the proportion of log-normally distributed excitatory connections within each pattern, and the proportion of log-normally distributed cross-pattern inhibitory connections. This allowed for a controlled comparison of network performance across different connectivity regimes (Fig. 5).

#### Synchrony is a more reliable predictor for familiarity detection across connectivity regimes

Rsync outperformed spike count in familiarity classification across all tested parameter combinations and exhibited consistently high performance across a broader range of connectivity regimes (Fig. 6A). This robustness to network connectivity is important. First, connectivity can change due to plasticity encoding new patterns in the excitatory population. Second, changing the level of competition between patterns modulates the density of inhibitory connections. Therefore, Rsync remains a reliable familiarity detection metric, even when the precise connectivity structure is variable or uncertain. In contrast, spike count-based familiarity detection fails when the number of encoded patterns increases. This is due to the overlap between patterns increasing, which increases connectivity between pattern clusters and results in higher output spike counts even when patterns are not familiar. In this regime, spike count was also more sensitive to the balance between excitatory and inhibitory connections, making it less reliable when connectivity is dynamic or noisy.

**Figure 6.**
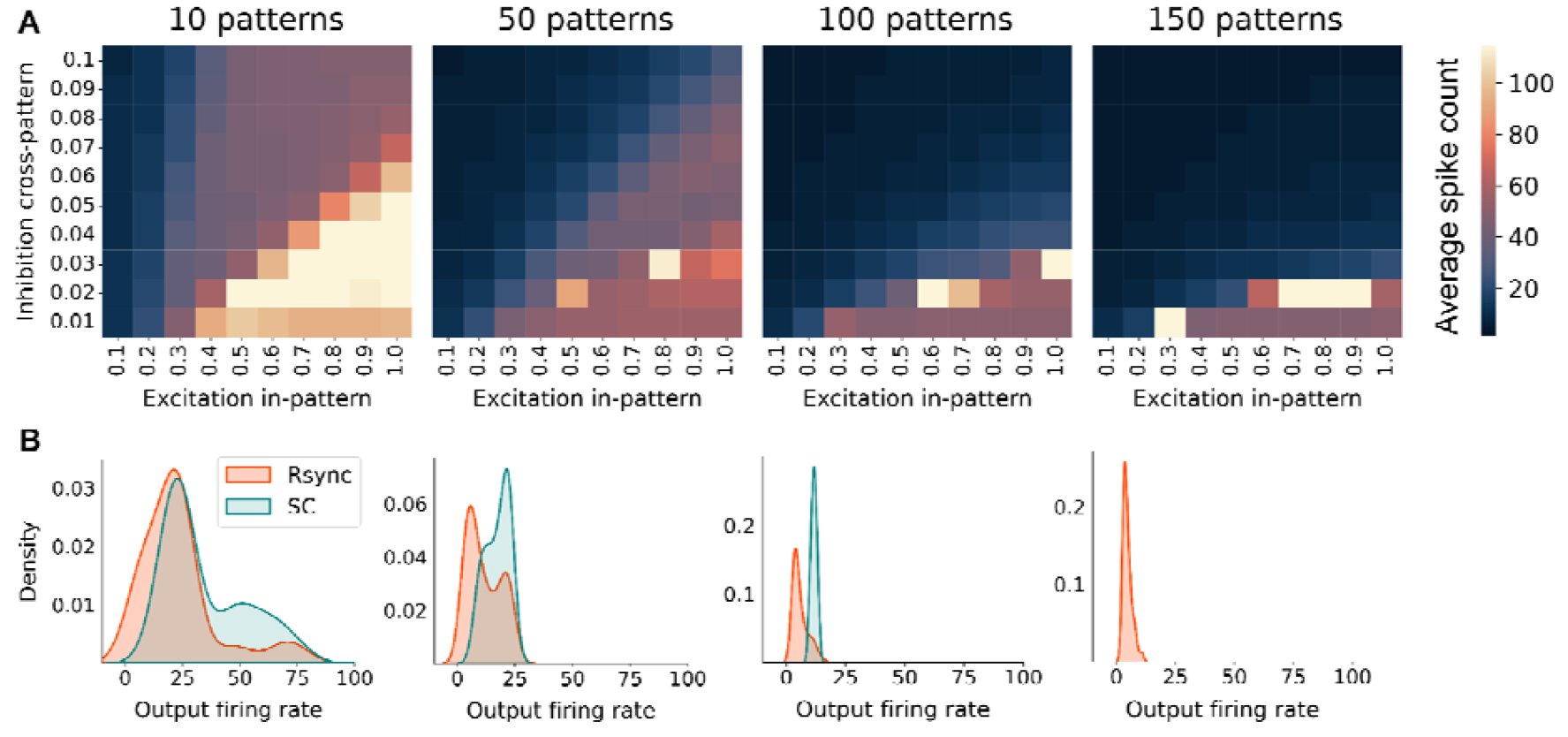
Network activity across connectivity parameters. Network activity is measured as the average output firing rate, for combinations of the proportion of in-pattern excitatory connections and cross-pattern inhibitory connections, with input saliency (firing rate) variability across samples 40 Hz. **A**. Average output firing rate. **B**. Distribution of output rate for parameter combinations that lead to F1 > 0.75 on the familiarity detection task. For 150 patterns, there is no parameter regime which would lead to this performance in a spike count familiarity detector. For networks with 10, 50, and 100 neurons, the distribution of Rsync values is significantly lower than that of spike count values (Mann-Whitney U test, *p* < 0.005 for all comparisons), showing that Rsync networks use spike-time codes and are more efficient.

#### Rate is a more reliable predictor for pattern classification across connectivity regimes

In contrast to its role in familiarity detection, spike count outperforms Rsync in classifying incoming patterns, regardless of the network’s connectivity structure (Fig. 5B). Rsync is less reliable in detecting new patterns, particularly when they overlap with familiar ones. This is because familiar patterns tend to evoke strong synchronous firing, reflecting the presence of established recurrent connections. New patterns, by contrast, typically have weaker within-pattern connectivity, leading to lower levels of recurrent input. Since new patterns are characterized by weak and scarce recurrent connectivity and in the presence of noisy input Rsync is driven by strong recurrent dynamics, its ability to distinguish between new patterns is limited. Spike count is directly influenced by feedforward drive, which means that it is better suited to identifying which specific pattern is currently active even when the pattern is unfamiliar. In summary, spike count more effectively classifies the input, while Rsync provides a stronger signal of how well the input matches the network’s learned connectivity.

#### Rate identifies neurons encoding a pattern, synchrony in that group encodes familiarity

Familiarity detection using synchrony is only reliable when measured locally, i.e. restricted to the neurons encoding the pattern in the network. Under a global decoding scheme, Rsync-based familiarity detection performance declined as the number of stored patterns increased (see Supplementary Fig. S2). Rate, on the other hand, reliably does identify the correct pattern encoding population, where synchrony can then be measured for familiarity detection. This underscores the benefit of combining rate and spike-time coding.

#### Optimal networks for complementary codes are balanced and sparse

Successful execution of both tasks − familiarity detection and pattern classification − depended on balanced excitation and inhibition at each neuron (BI Index avg. 0.04, see also Supplementary Fig. S1) [41]. At higher memory loads, this balance shifted slightly towards increased excitation and reduced inhibition. This adjustment compensates for greater between-pattern overlap, which increases cross-pattern inhibitory competition that would otherwise prevent persistent recurrent excitation necessary for successful encoding of familiarity. Individually, both Rsync and spike count performed best when total excitation and inhibition for every neuron was balanced (see also Supplementary Fig. S1).

Configurations with the best performance for familiarity detection from Rsync yield consistently significantly lower overall network activity compared to those performing best in spike count-based detection, regardless of memory load (Fig. 6). This matches the expectation that spike-timing codes can be more efficient, at least for familiarity detection, whereas spike count, encoding complementary information about the stimulus, will generally be less efficient. In our experiments, we consistently used rate coded poisson inputs. The efficiency of synchrony codes for familiarity forces the question if and when synchrony can also code for stimulus class, which we discuss next.

These results demonstrate that networks can simultaneously leverage both coding schemes with overlapping optimal parameter regimes. Importantly, these findings emerged despite using asynchronous Poisson inputs – an inefficient coding strategy due to sustained high firing rates and lack of coordinated temporal structure [9]. Nevertheless, our networks generated temporally coordinated output responses for familiar stimuli. Given this transformation capability from asynchronous to synchronous, we next examined how networks respond when inputs themselves possess temporal structure.

### From rate to synchrony coding

Our experiments showed that familiarity decoding under varied saliency relies on spike synchrony, whereas pattern classification depends primarily on rate coding. This suggests that stimulus content is propagated via firing rate, whereas spike synchrony conveys familiarity meta-information. This raises a natural question: Which coding strategy can networks employ once synchrony emerges through recurrent activity in response to familiar inputs? Hence, we repeated previous experiments on both familiarity detection and pattern classification, but increased synchrony of inputs (measured with Rsync, see methods for full experimental details)under the assumption that this input was synchronized in a preceding area.

#### Synchrony can detect stimulus class when temporal coordination of inputs increases

In our experiment, temporal synchronization of input patterns did not significantly affect familiarity detection performance: Rsync consistently outperformed spike count across a wide range of parameter configurations (see Supplementary Fig. S3). Similarly, spike count-based classification of patterns remained stable regardless of the degree of input synchrony (see Supplementary Fig. S4). However, Rsync-based pattern classification was sensitive to input synchronization: while it did not successfully classify fully decoherent Poisson inputs (see Fig. 5), it was able to classify patterns correctly when input synchrony increased (Fig. 7B-C). Under low memory loads − i.e., when fewer than 150 patterns were stored − Rsync reliably achieved high classification accuracy even with modest input synchrony. When inputs were perfectly synchronized, Rsync maintained high accuracy across nearly all parameter configurations.These findings suggest that, once neural activity synchronizes, temporal coherence can propagate through the sensory hierarchy, allowing both stimulus identity and familiarity to be decoded from spike synchrony alone.

**Figure 7.**
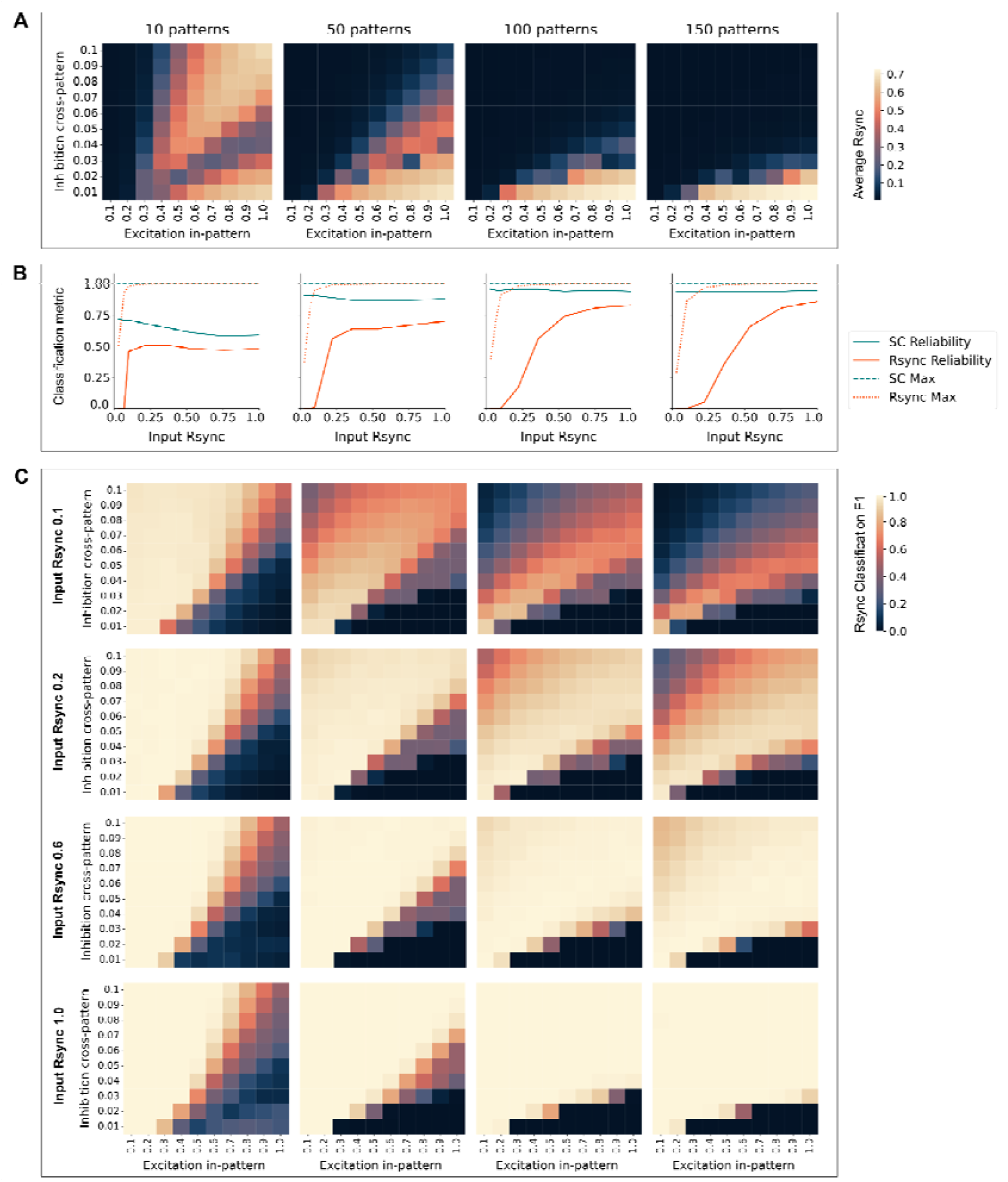
Temporal coordination of inputs enables reliable pattern classification with Rsync. **A**. Average Rsync values of inputs range from 0.08 to 0.75. **B**. Rsync and SC (spike count) reliability (the number of parameter configurations yielding F1 > 0.95) and maximal performance (maximum F1 across the configurations) for classification of patterns of different synchrony: 0.1, 0.2, 0.6, 1.0. Averaged over 10 trials, every trial includes 360 data samples, input saliency (firing rate) variability across samples 40 Hz. The more synchronized the inputs are, the more reliably Rsync can be used to classify them. **C**. F1 on Rsync-based pattern classification across the parameter configurations for input synchronization values: 0.1, 0.2, 0.6, 1.0.

**Fig 8.**
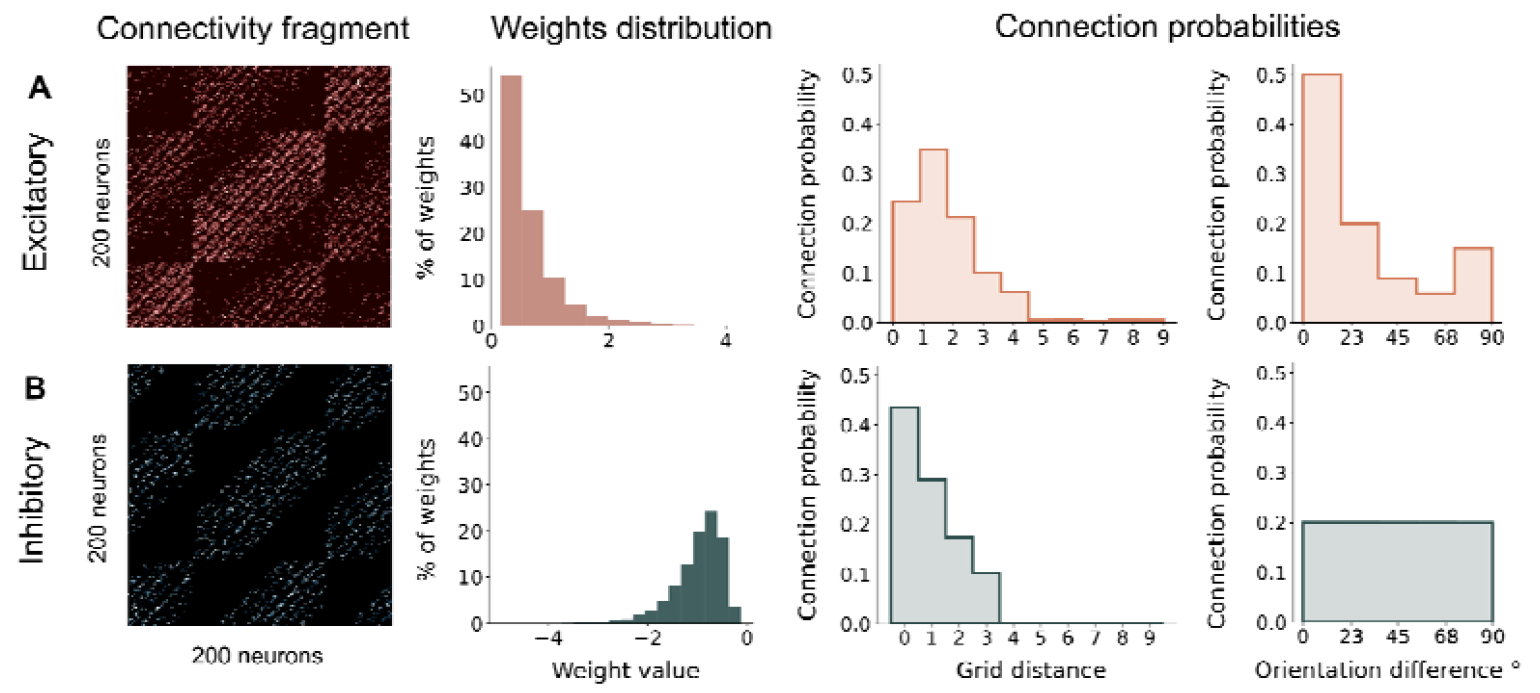
An example of V1 model connectivity. A fragment (200 neurons) of a heatmap, connection weights distribution, connection probabilities of the excitatory connectivity matrix of the: **A**. excitatory connectivity matrix, **B**. inhibitory connectivity matrix.

Synchrony appears to be an especially effective code for familiar patterns. When a pattern has been encountered before and induced Hebbian plasticity, strengthened recurrent connections lead to more synchronized responses, which Rsync can classify reliably. This synchrony can propagate through subsequent layers, allowing both class identity and familiarity to be decoded from the temporal structure of spike trains. In contrast, new patterns − activating neurons with weaker recurrent connectivity − produce more asynchronous responses, making spike count a more reliable signal. Thus, our results support the hypothesis that neural networks employ a complementary coding strategy: Familiar stimuli are encoded through efficient temporal codes, through synchrony while novel stimuli not yet encountered by the network are encoded through rate.

## Discussion

In this study, we have shown that synchrony and rate have complementary functions in a recurrent spiking neural network. Synchrony outperforms spike count as a measure of stimulus familiarity when input salience (modeled as varied input firing rate) changes, while spike count more reliably performs pattern classification. Both have important behavioral implications. Accurate pattern classification supports memory retrieval and generalization across contexts, while familiarity detection helps identify environmental changes, enables adaptation, and enables quick detection of known dangerous or rewarding stimuli. Our findings demonstrate that neural networks can simultaneously encode what a stimulus is and how familiar it is by leveraging distinct computational strategies, suggesting that the longstanding debate between rate and timing codes may be resolved through their functional complementarity.

The effectiveness of spike synchrony familiarity detection depends critically on where and how it is measured within the network. Our results show that spike synchrony responds more to recurrent connections than input rate, while output rate fails to separate these contributions. Synchrony is most effective locally, when measured from the subset of neurons receiving strong feedforward input. However, this creates an important limitation: identifying this specific group of neurons is difficult from synchrony alone, particularly for novel stimuli. Instead, spike count is required to reliably identify that neural population. This limitation directly arises from synchrony’s dependence on recurrent connectivity: novel inputs engage weakly connected neurons that haven’t yet formed a cluster, which then fails to generate synchronous activity necessary for reliable classification. At the same time, novel inputs that overlap with familiar input patterns sometimes recruit the population coding for the overlapped familiar stimulus, resulting in spurious synchrony that leads to false-positive classifications. In contrast, spike count is always primarily influenced by feedforward drive, making it a reliable predictor of stimulus identity regardless of its familiarity to the network. Previous work by our group [21] demonstrated that spike count can also be extremely efficient for relative familiarity detection of sequentially presented stimuli encoded with continuous plasticity. Under the more realistic condition of varied input saliency, this is no longer true. Here, our results align with and extend Brette’s theory of synchrony as a mechanism for invariant coding [42].

These mechanistic limitations change fundamentally when inputs are already synchronized rather than delivered as asynchronous Poisson spike trains. Under such conditions, network synchrony can reliably perform both familiarity detection and pattern classification functions. In this case, even weakly connected clusters coding for novel stimuli behave like strong ones; not by engaging recurrent connectivity, as familiar inputs do, but by achieving reliable activation through precise spike timing in the input. Thus, once activity synchronizes at a certain processing level, all information about the stimulus can be further propagated via synchrony coding. This result is consistent with anatomical and functional connectivity in the visual processing system in the brain. Early sensory areas like LGN are predominantly driven by vertical (feedforward or feedback) connections and lack lateral excitatory inputs [43], favouring rate-based representations. Conversely, higher cortical areas are increasingly densely connected laterally from V1 [44-45], though V2 [46] to IT [47]. This anatomical gradient suggests a natural functional transformation from purely rate-coded representations in early areas to increasingly synchrony and timing-based representations in higher-order cortical regions, in particular as stimuli become more familiar across that hierarchy.

In addition to a potential gradient along the sensory processing hierarchy, the complementarity of synchrony and rate codes also suggests complementary functional responses for processing familiar and novel stimuli. From the perspective of Bayesian modeling and predictive coding frameworks, familiar inputs match prior expectations, while novel inputs require prediction updates through synaptic plasticity [48]. Since recurrent connectivity can store prior experience in strongly connected clusters [19-20, 38], synchrony-based familiarity encoding likely supports completion and retrieval of familiar patterns by leveraging this encoded experience. This aligns with previous modeling work exploring phase-coded patterns as associative memory [49-50] and empirical evidence supporting the role of phase synchrony in memory formation and retrieval [51-52]. Conversely, strong novel stimuli induce strongly connected clusters through Hebbian plasticity in various network architectures [21, 37]. Importantly, this functional division also confers metabolic advantages: synchrony-based familiarity detection operates with lower overall firing activity, enabling more energy-efficient processing of familiar stimuli while reserving metabolically expensive rate-based plasticity for novel information that requires learning. Therefore, the idea of a complementary code for stimulus identity and familiarity also suggests a functional division: synchrony-mediated pattern completion for familiar stimuli versus rate-mediated plasticity induction for novel stimuli – a promising avenue for future investigations.

Understanding how our results relate to existing empirical work requires distinguishing between different types of familiarity. Most studies examine relative familiarity, the effect of recent exposure [53-54], whereas we model lifelong absolute familiarity as the match between stimulus patterns and learned recurrent connectivity. Our models exhibit increased activity for familiar stimuli, contrasting with empirical findings on relative familiarity recorded in multiple cortical areas, including V2, the inferotemporal cortex (IT), the perirhinal cortex (PrC), and the prefrontal cortex, where familiar stimuli lead to repetition suppression [55-57]. However, multiple studies have reported that the electrophysiological markers of lifelong absolute familiarity are distinct from those of relative familiarity [58-59], which suggests a different underlying mechanism and explains the different activity patterns. Consistent with this distinction, a recent V1 study showed that phase-reversing familiar visual stimuli elicit an initial increase in V1 firing rate, followed by prolonged suppression [60], suggesting that familiarity-induced excitation can precede strong inhibition. Thus, the increased activity in response to familiar stimuli in our model may also be attributed to our simple implementation of inhibition that lacks explicit dynamic interactions between excitatory and inhibitory neurons. Future work could therefore explore more detailed inhibitory circuitry, which could enable repetition suppression caused by complex inhibitory circuits.

The complementary coding strategy suggested here optimizes both efficiency and robustness in neural computation. Spike synchrony excels at familiarity detection and operates efficiently with few spikes required across varying stimulus saliency levels, but it has a critical limitation: it consistently misclassifies novel stimuli because synchrony depends on established recurrent connections that novel patterns lack. Firing rate coding solves this problem by providing robust stimulus classification regardless of connectivity strength. This suggested that we can select network configurations that optimize for both. The result is a division of labor where synchrony efficiently handles familiarity detection while rate coding ensures robust classification of all stimuli − both familiar and novel − within a single, energy-efficient network architecture. This complementary relationship thus provides a mechanistic framework for how neural circuits can simultaneously achieve computational efficiency and the flexibility needed to process dynamic sensory environments.

## Methods

### Familiarity detection

To classify stimulus familiarity, we extracted group-level features from simulated spike trains: a rate-based (Eq. 1) and a synchrony-based measure capturing temporal dynamics (Eq. 2).

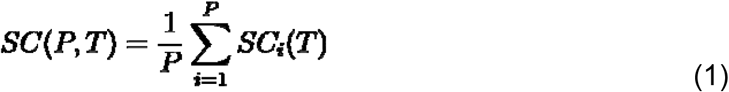

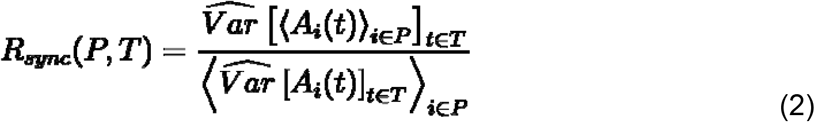

Here,***P*** defines the externally-stimulated neuron population representing one encoded pattern, and ***T* = 500** ms is a time interval for measuring spike count or synchrony. Eq. 1 describes population spike count ***SC***, which is measured as an average spike count of all neurons in the population representing one pattern.

In Eq. 2, Rsync measures spike synchrony by comparing the variance of the population mean activity to the average variance of individual neurons. ***A***_***i***_ is an activation trace of neuron ***i*** from neural population ***P***. To compute activation traces, raw binary spike trains were first convolved with an exponential kernel ***k*(*t*) *= exp* (*−t/τ*)**, with timescale ***τ* = 5** ms. If all neurons fire in perfect synchrony, their activity traces are identical, so the variance of the mean field equals that of individual neurons, yielding ***Rsync* = 1;**. Likewise, if neurons fire independently, the mean field gets averaged out and flattened, and ***Rsync* → 0**. Unlike pairwise correlations, ***Rsync*** is a group measure which scales to populations of any size.

For each model, we defined network connectivity structure, selected neurons for performance measurements and reported F1 scores obtained through 10-fold cross-validation. We also show baseline scores for random predictions.

### Pattern classification

In addition to familiarity detection, we performed multi-class pattern classification to identify which stimulus pattern the network was currently representing. Each dataset contained 160 novel and 160 familiar patterns, resulting in 320 classification instances per run. For this task, classification was independent of familiarity: the goal was to recover the identity of the stimulated population of neurons from the output activity.

We applied a simple maximum-response rule using both rate-based (spike count, Eq. 1) and synchrony-based (Rsync, Eq. 2) metrics. Spike count and synchrony were computed separately for each candidate neural population. The population yielding the maximal value of either measure was taken as the classification result − decoded pattern identity. This winner-take-all procedure produces a one-of-N classification, where N is the total number of new and familiar predefined patterns. Once the population was classified, i.e. selected by the max-rule, familiarity detection was performed only for that population, as described in the previous section.

### Spiking model

In our experiments, we used spiking recurrent networks (SNNs) composed of Leaky Integrate- and-Fire (LIF) neurons. The membrane potential of neurons evolves according to the equation

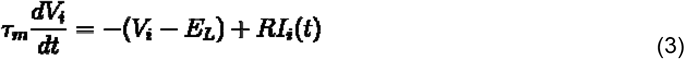

Here, ***dV*** is an update of membrane potential per timestep ***dt* = 1.0** ms, resting potential ***E***_***L***_ **= −65** mV, membrane time constant ***τ* = 10** ms. The firing threshold in the model is set to −55 mV, and the refractory period equals 3 ms. ***I***_***i***_**(*t*)** are the synaptic currents arriving at the soma:

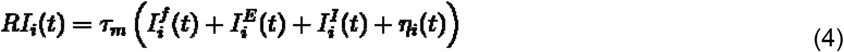

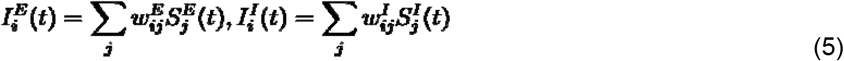

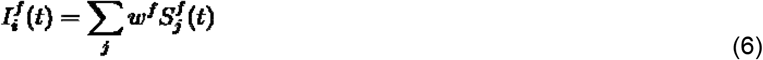

These currents include summed input from feedforward, recurrent excitatory and recurrent connections, scaled by corresponding PSP efficacy values ***w***^***f***^,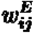 and 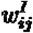. ***S***_***j***_**(*t*)** denotes a presynaptic spike arrival as the Dirac delta function ***S***_***j***_**(*t*) = *δ* (*t − t***_***j***_). All feedforward weights are equal and set to 3. We did not simulate separate excitatory and inhibitory neuron populations. Instead, each neuron projects both excitatory and inhibitory outputs, defined by two distinct recurrent connectivity matrices. External input is delivered via one-to-one feedforward connections, modeled as a Poisson process.

Additionally, we model random fluctuations of the membrane potential due to stimulus independent background activity by including the Gaussian white noise term **⟨ *η***_***i***_ **(*t*) ⟩** equal 0.6 with ***σ***^**2**^ equal 0.4 mV. The membrane noise is uncorrelated across neurons and time **⟨ *η***_***i***_ **(*t*) ⟩** and ***σ***^**2**^ biases neurons towards firing so that they can be driven effectively by very sparse Poisson inputs within our 500 ms simulation time. We chose maximal and values under which neurons only fire upon receiving external input, despite the noise. Thus, neurons’ membrane potentials were consistently kept in a subthreshold state.

### V1 model connectivity

Each neuron in the model has a receptive field defined by spatial and orientation selectivity. The model is designed to recognize 8 orientation angles (from 0° to 180° in 22.5° steps) using 800 neurons, each tuned to a specific orientation. These neurons are arranged in a 10×10 grid representing a visual field. We measure distance between neurons as Chebyshev distance − the maximum difference along any coordinate dimension, appropriate for a 2D grid setup. Each unit of difference corresponds to 100 µm.

Excitatory connections exhibit 10% sparseness and follow a log-normal weight distribution, consistent with measurements of pyramidal neuron connectivity in layers 2/3 of V1 [27, 61]. Connection probability depends on both spatial proximity and receptive field similarity. While most connections form over short distances, the model also includes a small number of strong long-range recurrent connections, in line with empirical observations [62].

Inhibitory connections are modeled after Parvalbumin expressing (PV) interneurons – the prevalent type of inhibitory interneurons in layers 2/3 of V1. These neurons exhibit weak orientation selectivity [63], in our model simplified to no orientation selectivity, as in Taylor et al. [64]. The inhibitory connectivity depends on distance, follows log-normal distribution [61, 65] and has a total sparseness of 6% (20% within 300 µm). As a result, our V1 model has a broad, non-selective inhibitory mechanism that serves primarily to regulate overall excitatory activity, which could otherwise destabilize the recurrent network.

### Synchronized input generation

To control the level of input synchronization, we generated patterns using the following procedure: (1) identical Poisson spike trains were first generated and assigned to all neurons within a given pattern; (2) all spike events were then jittered independently across all stimulated neurons within a specified time window. By varying the width of this window, we systematically modulated the resulting input synchrony (Table 1), measured as Rsync (see Eq 2).

**Table 1.** Input synchrony. Temporal jitter window values used to generate patterns with specific levels of input Rsync, averaged over 100 trials for each jitter value.

|  |  |
| --- | --- |
| $1.0 \pm 0.0$ | 0 |
| $0.55 \pm 0.04$ | 5 |
| $0.2 \pm 0.06$ | 20 |
| $0.1 \pm 0.03$ | 50 |

## Data and code availability

All original data in the work was programmatically generated. The code for data generation, analysis and visualization of the presented results can be found at https://github.com/rainsummer613/snn-saliency-familiarity-coding.

## Acknowledgments

We thank the Deutsche Forschungsgemeinschaft (DFG, German Research Foundation) for funding the Priority Programme SPP 2262 *Memristive Devices Toward Smart Technical Systems* (422738993). The high-performance computing cluster used for the simulations was also funded by the DFG (456666331) and the Ministry for Science and Culture of Lower Saxony (MWK). The funders had no role in study design, data collection and analysis, decision to publish, or preparation of the manuscript.

## Author Contributions

V.Z. and P.N. conceived and designed the study. V.Z. implemented the computational models, conducted analyses, and prepared all visualizations. P.N. and G.P. supervised the research and provided resources. V.Z. and P.N. wrote the initial draft, and all authors contributed to review and editing of the manuscript. Funding was acquired by G.P.

