## Supplementary materials for "When Firing Rate Falls Short: Spike Synchrony Reliably Disentangles Stimulus Saliency and Familiarity"

### Fig S1

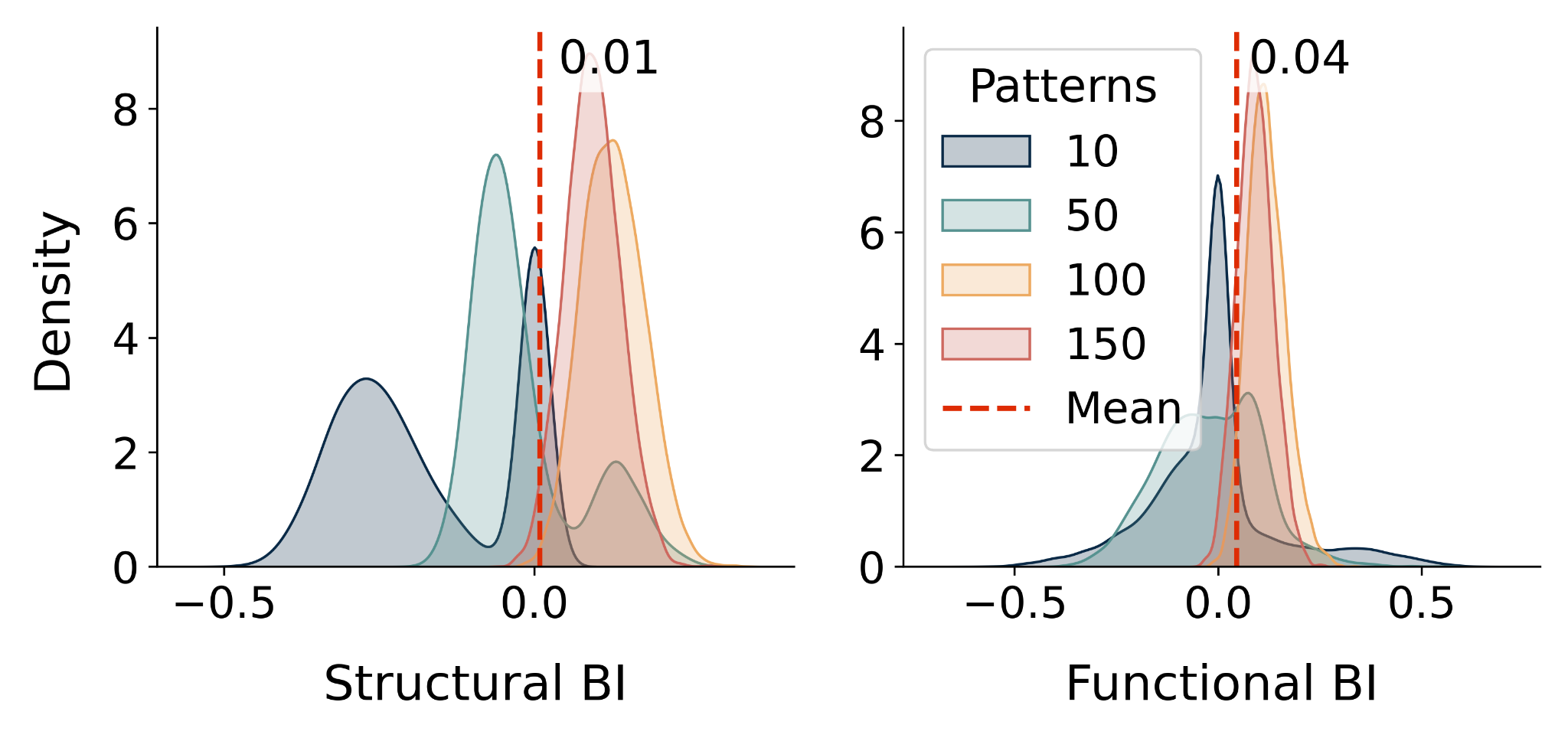

**Fig S1. Optimal E/I balance for familiarity detection**

Excitation ratio for the 5 best-performing configurations of in-pattern excitatory and cross-pattern inhibitory connectivity, across different memory loads (10, 50, 100, 150 patterns). **A**. Balance Index (BI) for weighted incoming connections per neuron, computed as the average normalized difference between the weighted sums of incoming excitatory and inhibitory weighted connections. **B**. Balance Index (BI) for total lateral input per neuron, computed as the average normalized difference between excitatory and inhibitory input across time steps.

### Fig S2

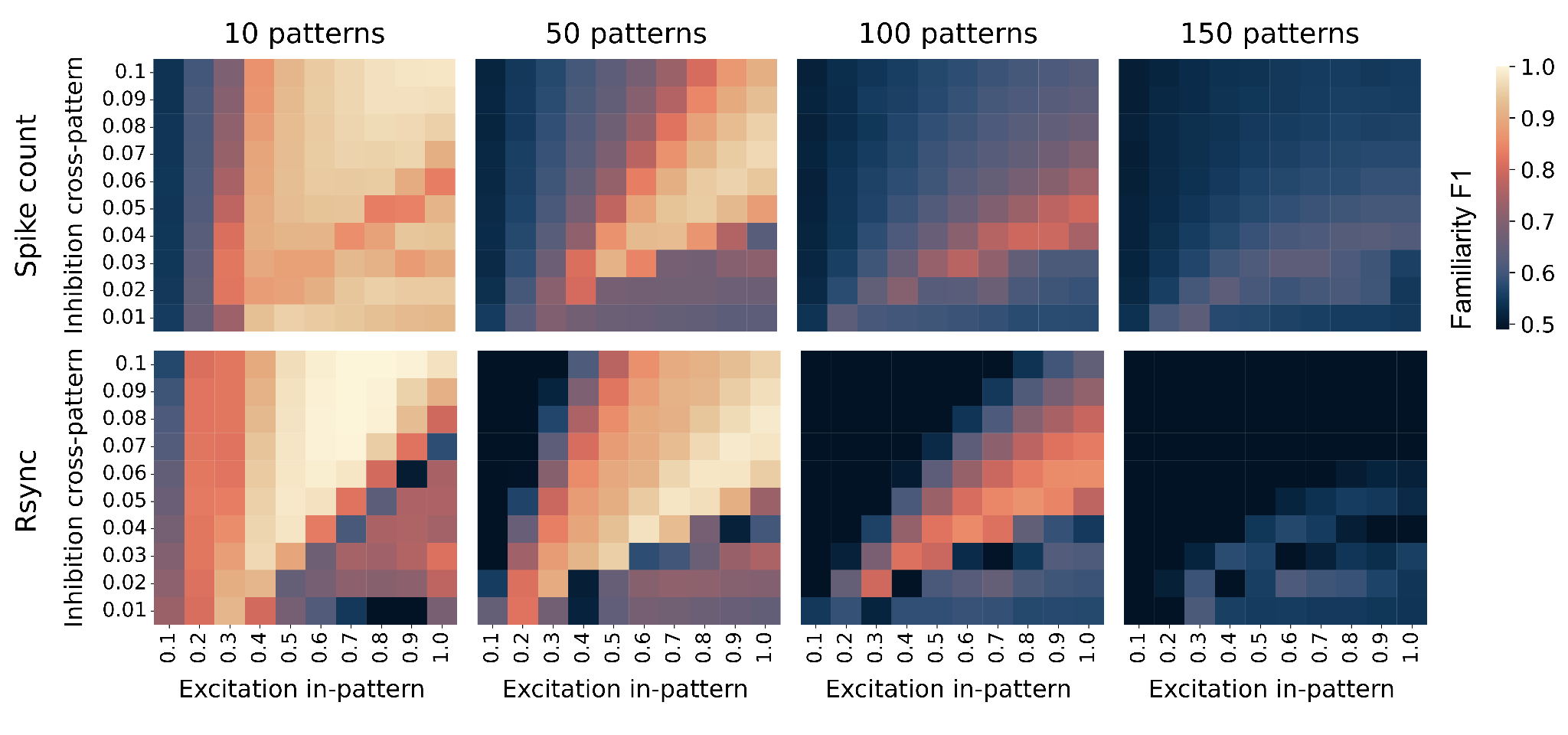

**Fig S2**. **Familiarity detection from globally measured Rsync and spike count**

Cross-validated performance of global, i.e. measured across all neurons, Rsync- and spike count-based classifiers, for every combination of the proportion of in-pattern excitatory connections and cross-pattern inhibitory connections. Averaged over 10 trials, every trial includes 360 data samples, input saliency (firing rate) variability across samples 40 Hz. Both metrics’ performance drop rapidly with the increasing number of patterns.

### Fig S3

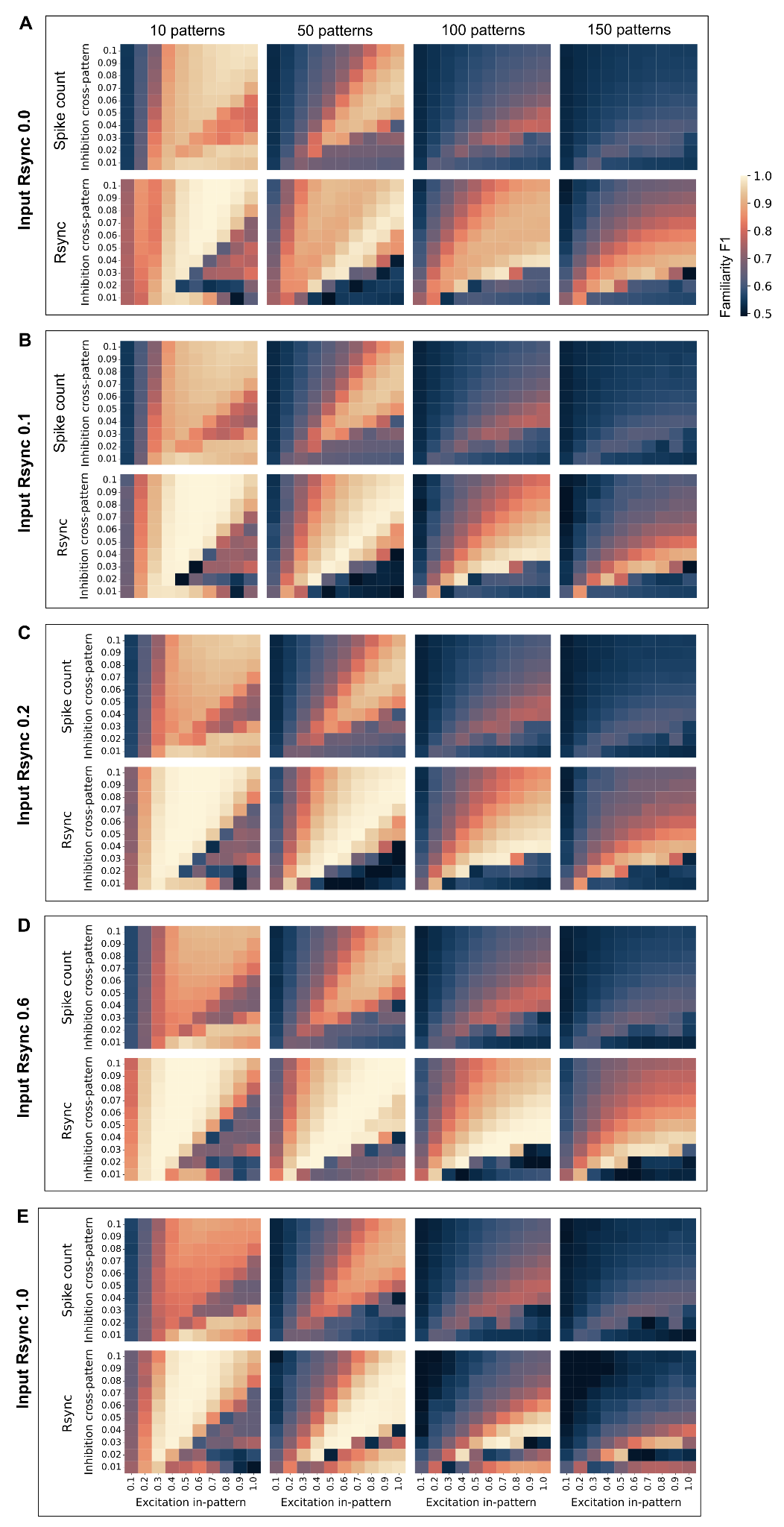

**Fig S3**. **Familiarity detection of synchronized inputs**

Cross-validated performance (F1 score) of Rsync- and spike count-based familiarity detectors (binary classifiers), for every combination of the proportion of in-pattern excitatory connections and cross-pattern inhibitory connections, compared across different levels of input synchrony measured as Rsync (see Eq 2 and Methods). Averaged over 10 trials, every trial includes 360 data samples, input saliency (firing rate) variability across samples 40 Hz. Rsync consistently outperforms spike count across most parameter configurations regardless of the input synchronization level.

### Fig S4

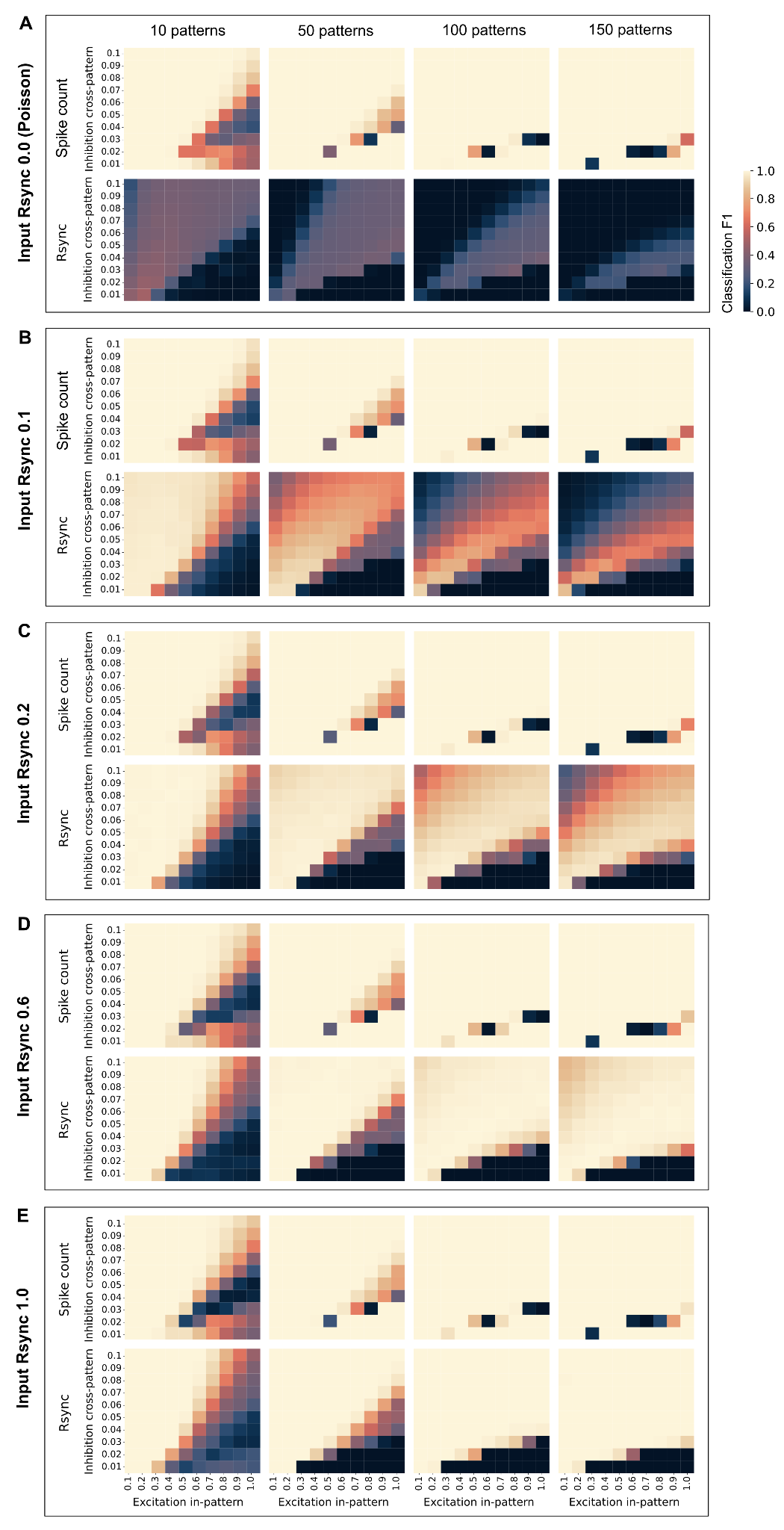

**Fig S4**. **Classification of synchronized inputs**

Cross-validated performance (macro F1 score) of Rsync- and spike count-based familiarity detectors, for every combination of the proportion of in-pattern excitatory connections and cross-pattern inhibitory connections, compared across different levels of input synchrony measured as Rsync (see Eq 2 and Methods). Averaged over 10 trials, every trial includes 360 data samples, input saliency (firing rate) variability across samples 40 Hz. Spike count consistently performs excellent regardless of the input synchronization level, while Rsync performance gets better with increasing input synchronization. Highly synchronous inputs can be very reliably classified by both spike count and Rsync.
